# Trophic transfer of microplastics enhances plastic additive accumulation in fish

**DOI:** 10.1101/2021.03.09.434685

**Authors:** Takaaki Hasegawa, Kaoruko Mizukawa, Bee Geok Yeo, Tomonori Sekioka, Hideshige Takada, Masahiro Nakaoka

**Affiliations:** Graduate School of Environmental Science, Hokkaido University; Akkeshi, Hokkaido, Japan; Laboratory of Organic Geochemistry, Tokyo University of Agriculture and Technology; Fuchu, Tokyo, Japan; Faculty of Fisheries Sciences, Hokkaido University; Hakodate, Hokkaido, Japan; Akkeshi Marine Station, Field Science Center for Northern Biosphere, Hokkaido University; Akkeshi, Hokkaido, Japan

## Abstract

Organisms ingest microplastics directly from environments and indirectly from food sources. Ingesting microplastics can lead to an organism accumulating plastic-derived chemicals. However, the relative contributions of the two exposure routes (direct ingestion vs. indirect ingestion) to plastic-derived chemical accumulation in an organism are unknown. Using microplastics containing different types of plastic additives, we performed exposure experiments to compare chemical accumulation patterns in fish (*Myoxocephalus brandti*) between the exposure from the water and prey (*Neomysis* spp.). We found significantly higher brominated flame retardant concentrations in fish fed microplastic-contaminated prey than fish exposed to microplastics suspended in the water. The results indicate that prey-mediated ingestion of microplastics can be a more significant route for organisms accumulating plastic-derived chemicals, suggesting that organisms at higher trophic levels are more exposed to microplastics and associated chemicals than organisms at lower trophic levels.

## Introduction

Ingestion of microplastics by marine organisms has become a common phenomenon in global oceans, raising concerns about the impacts on marine ecosystems [1]. Marine predators are exposed to microplastics by ingesting them directly from the water column and indirectly from microplastic-contaminated prey [2]. Recent studies have shown that fish can ingest larger amounts of microplastics in prey than from the water column [3–5]. Microplastics can negatively affect organisms that consume them [6]. Prey-mediated ingestion of microplastics by predatory organisms is therefore of great concern.

Plastics contain various chemical additives that are incorporated during manufacturing [7]. It has been found that ingesting plastics can result in the chemical additives in the plastic accumulating in marine organisms [8–10]. Larger amounts of microplastics can be ingested by fish in prey than directly from the water column, so consuming plastic-contaminated prey can greatly increase the amounts of various chemicals contained in the plastics that accumulate in predatory fish.

The aim of this study was to assess the relative contributions of direct and indirect exposure to microplastics (exposure to plastics in the water versus exposure to plastics in prey, respectively) on accumulation of chemical additives in predatory fish. We used the same model prey–predator system (the mysids *Neomysis* spp. and the benthic fish *Myoxocephalus brandti*) as was used in a previous study [4]. We performed exposure experiments using polyethylene microplastics containing two brominated flame retardants (BFRs) and three ultraviolet stabilizers (UVSs). The relative importances of direct and indirect exposure were assessed by exposing fish to polyethylene microplastics suspended in the water column (for direct, waterborne exposure) or to mysids previously exposed to polyethylene microplastics (for indirect, prey-mediated exposure). We then determined the concentrations of the additives in the muscle and liver tissues of the fish and of fish collected immediately after the fish were collected from the field (ambient control samples).

## Materials & Methods

### Animal collection and acclimation

Procedures for capture and handling of animals conformed to the Committee of Animal Ethics in the Hokkaido University (31-4).

We collected organisms for use in the experiment on 7 September 2020 from a seagrass bed in Akkeshi-ko estuary (43°02′ N, 144°52′ E) off eastern Hokkaido, northern Japan. Mysids (*Neomysis* spp., mean ± standard deviation body length 7.70 ± 1.53 mm, mean ± standard deviation wet weight 8.20 ± 3.59 mg) and juvenile *M. brandti* (mean ± standard deviation body length 7.40 ± 0.52 cm, mean ± standard deviation wet weight 4.35 ± 0.81 g) were collected using an epibenthic sled. The concentrations of the contaminants of interest before the experiment were determined by analyzing 90 mysids and three fish that were frozen immediately after being collected. Living mysids and fish were also transported to the laboratory for use in the experiments. The mysids were acclimated in a number of 30 L aquaria overnight with flowing seawater that had been filtered through fine sand to allow potential microplastics to be cleared from the guts of the mysids. Ten glass aquaria (each 245 mm × 165 mm × 160 mm) were prepared for use in the experiment. One randomly selected fish was placed in each aquarium and allowed to acclimate for 7 d before the experiment started. During the acclimation period, the mysids were fed microalgae (Shellfish diet 1800; Reed Mariculture) each day and the fish were fed fresh mysids.

### Preparation of plastics with additives

Cylindrical low-density polyethylene pellets (diameter 5 mm, length 5 mm; DJK Corporation, Chiba, Japan) containing five additives were used in the experiment. Polyethylene is one of the most common plastic polymers and is commonly found in the oceans [7]. The five additives were added to the polyethylene pellets during the pellet production process. The additives were 1,2,3,4,5-pentabromo-6-(2,3,4,5,6-pentabromophenoxy)benzene (BDE-209, CAS no. 1163-19-5), 1,2,3,4,5-pentabromo-6-[2-(2,3,4,5,6-pentabromophenyl)ethyl]benzene (DBDPE, CAS no. 84852-53-9), 2-(benzotriazol-2-yl)-4,6-bis(2-phenylpropan-2-yl)phenol (UV-234, CAS no.70321-86-7), 2,4-ditert-butyl-6-(5-chlorobenzotriazol-2-yl)phenol (UV-327, CAS no. 3864-99-1), and (2-hydroxy-4-octoxyphenyl)-phenylmethanone (BP-12, CAS no. 1843-05-6). BDE-209 and DBDPE are widely used as flame retardants in plastic products and electrical appliances and are suspected not to be readily degradable and to have long-term toxic effects [11]. UV-234, UV-324, and BP-12 are commonly added to plastic products to act as UV stabilizers. UV-234, UV-324, and BP-12 have the potential to bioaccumulate and cause toxic effects in organisms [12]. These compounds were selected because they have previously frequently been detected in plastic debris collected from natural environments and in marine organisms [12–14]. The concentration of each additive in the polyethylene pellets was <0.4% by weight to be in the same order of magnitude as concentrations found in plastic debris collected from natural environments and in marine organisms. Polyethylene powder (Flo-Thene, FG701N; Sumitomo Seika Chemicals Co., Osaka, Japan) and authentic standards of the additives in powder form (Wako Pure Chemical Industries, Osaka, Japan) were mixed well and molded into pellets in a co-rotating twin-screw kneading extruder (HK-25D; Parker Corporation, Tokyo, Japan). The pellets were melted and re-extruded twice to ensure that the additives were uniformly distributed within the pellets. The pellets were then ground to powder using a plastic cutting mill (PLC-2M; Osaka Chemical Co., Osaka, Japan), then the powder was passed through 63 and 32 μm mesh sieves. The particles remaining on the 32 μm sieve were used in the experiment.

The particle sizes were determined using a quantification method using Nile red and a nylon mesh filter with 8 μm pores [15]. The particle area was measured, and the square root of the area was defined as the particle size. The polyethylene microplastic particle size was 29.90 ± 13.40 μm (mean ± standard deviation), and the size distribution is shown in Fig S1. The concentrations of the five additives in the polyethylene microplastics were determined and are shown in Table 1, and the contributions of the additives to the total additive concentration are shown in Fig S2.

**Table 1.**
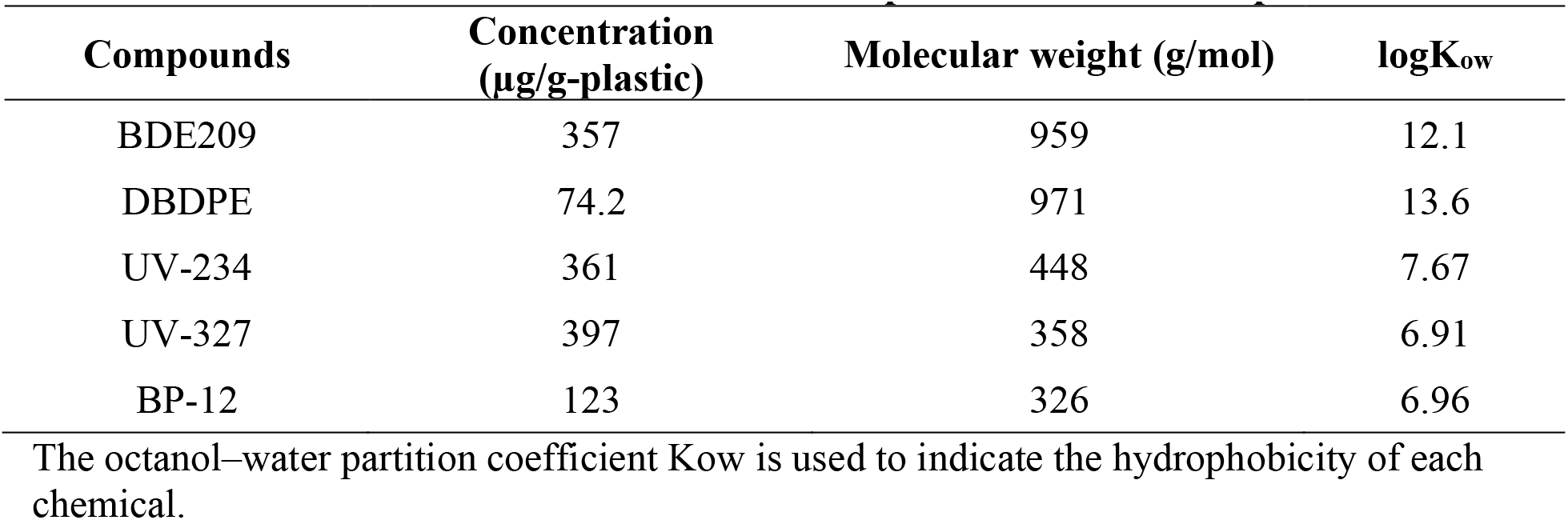
Information on the additives in the microplastics used in the experiment.

| **Compounds** | **Concentration**<br>**($\mu\text{g/g-plastic}$)** | **Molecular weight (g/mol)** | **logK<sub>ow</sub>** |
| --- | --- | --- | --- |
| BDE209 | 357 | 959 | 12.1 |
| DBDPE | 74.2 | 971 | 13.6 |
| UV-234 | 361 | 448 | 7.67 |
| UV-327 | 397 | 358 | 6.91 |
| BP-12 | 123 | 326 | 6.96 |

The octanol–water partition coefficient Kow is used to indicate the hydrophobicity of each chemical.

To prepare a microplastic suspension, 100 mL of distilled water in a glass beaker was brought to the boil in a microwave oven, then 10 μL of surfactant (Tween 80; polyethylene sorbitol ester; Cospheric LLC) were added and the solution was stirred with a glass rod for 30s. A 5 mL aliquot of the solution was then transferred to another glass beaker containing 95 mL of boiled distilled water to give a 0.005% surfactant solution. The solution was left at room temperature for 1 h. Before the exposure experiment was started, 8 mg of polyethylene microplastics were added to a plastic tube containing 8 mL of the 0.005% surfactant solution. The tube was vortexed and left at room temperature for 1 h to ensure that the polyethylene particles were immersed in the solution. A fresh microplastic suspension was prepared for each experiment to minimize leaching of chemicals from the polyethylene microplastics during the immersion process.

### Leaching experiment

The amounts of the chemicals that leached from the microplastics into seawater were determined using three beakers each containing 1 L of filtered seawater and microplastics at a concentration of 2 mg/L. The mixtures were prepared and left for 24 h with the seawater constantly circulated and aerated. At the end of the 24 h period, the seawater was transferred to a 1 L stainless steel container and stored in a freezer until it was analyzed.

### Ingestion experiment for mysids

A total of 30 1 L glass beakers were prepared and divided into two groups. To each of the 15 bottles in one group were added 1 L of seawater that had been passed through a 1 μm filter and microplastics to give a concentration of 500 μg/L. The microplastic concentration was in the same order of magnitude as the microplastic concentration of 250 μg/L previously found in water in one of the most heavily contaminated areas in the world [16]. To each of the 15 bottles in the other group was added 1 L of leachate seawater. The leachate seawater was prepared by adding microplastics at a concentration of 500 μg/L to 1 L of seawater that had been passed through a 1 μm filter and then passing the suspension through a filter to remove the microplastics. The leachate seawater was used to assess uptake by the mysids of chemicals leached from the microplastics into the seawater during the experiment. Mysids in the acclimation aquaria were randomly selected and three were placed in each bottle. The mysids were fed 2 mg (dry weight) of microalgae. The seawater was constantly circulated and aerated to ensure that the microplastics were evenly distributed in the bottles. After 24 h, the mysids were flushed gently with filtered seawater to remove beads from their exoskeletons and their body lengths and wet weights were determined. The mysids were then frozen and stored at −30 °C until they were analyzed. The same procedure was repeated for 4 d, and 300 mysids in total were collected from each group except that only 30 mysids were used in each group on the fourth day.

### Trophic transfer experiment

After the fish had acclimated, 10 aquaria were prepared and divided into two groups. To each of five aquaria were added 2 L of seawater that had been passed through a 1 μm filter, microplastics to give a concentration of 500 μg/L, and nine plastic-free mysids (meaning the fish would ingest microplastics only from the water column). To the other five aquaria were added nine mysids that had previously been exposed to microplastics in seawater at a concentration of 500 μg/L and 2 L of seawater that had been passed through a 1 μm filter. No microplastics were added to ensure that the fish only consumed microplastics in the mysids. The number of mysids was determined from the results of a preliminary experiment to ensure that all of the mysids were consumed by the fish within 24 h. The mysids were exposed to microplastics following the procedure used in the first experiment described above. The exposure experiment lasted 10 d. The seawater in each aquarium was exchanged once each day. The seawater was constantly circulated and aerated throughout the experiment to ensure that the microplastics were evenly distributed. We did not observe any microplastics on the bottoms of the aquaria throughout the experiment. After 10 d of exposure, the fish were gently flushed with filtered seawater to remove beads from the body surfaces. The body lengths and wet weights of the fish were determined, then the fish were frozen and stored at −30 °C until they were analyzed.

### Sample analysis

#### Chemicals and reagents used in the analyses

Acetone, n-hexane, 2,2,4-trimethylpentane (iso-octane), methanol, pyridine, acetic anhydride, hydrochloric acid, anhydrous sodium sulfate, Wakogel Q-22 (75 μm, 200 mesh), and Wakogel Q-23 (75–150 μm, 100–200 mesh) were purchased from Wako Pure Chemical Industries. Dichloromethane (DCM) was purchased from Kanto Chemical Co. (Tokyo, Japan). BDE-209 and DBDPE analytical standards were purchased from Wellington Laboratories (Guelph, Canada). Individual hexa- to nona-chlorinated diphenyl ether congeners were obtained from Wellington Laboratories (BDEs-179, −183, −188, and −202), Cambridge Isotope Laboratories (Tewksbury, MA, USA) (BDE-207), and Accustandard (New Haven, CT, USA) (BDEs-155, −184, −196, −197, −203, −206, and −208). UV-327 and BP-12 were obtained from Accustandard. The surrogate standards were f-BDE-208 (4′-fluoro-2,2′,3,3′,4,5,5′,6,6′-nonabromodiphenylether) supplied by Chiron AS (Trondheim, Norway) for the BFRs, UV-327-d20 supplied by Toronto Research Chemicals (North York, Canada) for UV-327 and UV-234, and BP-12-d17 supplied by Hayashi Pure Chemical Ind. (Osaka, Japan) for BP-12. The internal injection standard, chrysene-d12, was purchased from Sigma-Aldrich (St. Louis, MO, USA). All glassware was rinsed with methanol, acetone, and distilled n-hexane three times each or baked at 550 °C for 4 h before use.

#### Biological sample analysis

The livers and whole body tissues (mainly the muscle tissues, excluding the intestines, head, and fins) of the fish were analyzed. Each sample was freeze-dried and then extracted using an ASE200 accelerated solvent extractor (Dionex, Sunnyvale, CA, USA) using a 1:3 v/v mixture of acetone and DCM. Each sample was extracted in an 11 mL stainless steel cell using a preheating time of 0 min, a heating time of 5 min, a static time of 5 min, a flush of 100%, a 60 s purge, two cycles, a pressure of 1500 psi, and a temperature of 100 °C.

An aliquot of each whole-body sample extract was transferred to a weighed 4 mL vial and evaporated to dryness to determine the lipid content of the sample. The surrogate standards (f-BDE-208, UV-327-d20, and BP-12-d17) were added to another aliquot of the extract, then the extract was evaporated just to dryness in a rotary evaporator and transferred to a 10 mL centrifuge tube. The extract was evaporated to <100 mL under a gentle stream of nitrogen, then 50 mL each of pyridine and acetic anhydride were added. The mixture was kept at room temperature for >8 h, then the reaction was stopped by adding 200 mL of 4 M HCl. Acetylates were then extracted with n-hexane five times. The n-hexane extract was passed through anhydrous sodium sulfate to remove water and collected in a pear-shaped flask.

The solvent was exchanged from n-hexane to DCM, then the acetylated sample was purified by gel permeation chromatography. About 2 mL of an extract was passed through a gel permeation chromatography column (2 cm i.d., 30 cm long; CLNpak EV-2000; Showa-denko, Tokyo, Japan) to separate the target compounds from biolipids. The mobile phase was DCM and the flow rate was 4 mL/min. The fraction with a retention time of 12–25 min was collected and purified further.

After gel permeation chromatography, the extract was evaporated just to dryness in a rotary evaporator and then passed through a column (1 cm i.d., 9 cm long) packed with silica gel (Wakogel Q-22) that had been deactivated by adding 10% w/w H2O. The first fraction, containing BFRs, was eluted with 35 mL of a 3:1 v/v mixture of n-hexane and DCM. The second and third fractions, containing UVSs, were eluted with 15 mL of a 1:99 v/v mixture of methanol and DCM and 20 mL of a 1:4 v/v mixture of 1% methanol in DCM and acetone, respectively. The second and third fractions were combined, evaporated to a small volume, and transferred to a 1 mL amber ampoule for instrumental analysis.

The first fraction, containing BFRs, was evaporated just to dryness using a rotary evaporator and then passed through a column (0.47 cm i.d., 18 cm long) containing fully activated silica gel (Wakogel Q-23). The first fraction eluted from the column, containing aliphatic hydrocarbons, was eluted with 5 mL of n-hexane. The second fraction, containing BFRs, was eluted with 15 mL of an 85:15 v/v mixture of n-hexane and DCM or 10 mL of a 3:1 v/v mixture of n-hexane and DCM. The BFR fraction was evaporated to a small volume and transferred to a 1 mL amber ampoule for instrumental analyses.

#### Water sample analysis

Each water sample from the leaching experiment was passed through a glass fiber filter (GF/F). The surrogate standards f-BDE-208, UV-327-d20, and BP-12-d17 dissolved in acetone were then added to the filtrate. The solution was then liquid–liquid extracted with DCM (10% of the water sample volume), and the DCM was dried using anhydrous sodium sulfate. The liquid–liquid extraction process was repeated three times. The extracts were combined, evaporated using a rotary evaporator, and then derivatized, fractionated, and purified using the acetylation and 10% water-deactivated silica gel (Wakogel Q-22) column chromatography methods described above for the biological samples.

#### Instrumental analysis

For BFR analysis, a sample was evaporated just to dryness under a gentle stream of nitrogen and then the residue was redissolved in 100 mL of iso-octane.

The UVSs were determined using a 7890 gas chromatograph and 5977 quadrupole mass spectrometer (Agilent Technologies, Santa Clara, CA, US). Separation was achieved using an HP-5MS 30-m fused silica capillary column (0.25 mm i.d., 0.25 mm film thickness; Agilent Technologies). The carrier gas was helium, and the carrier gas pressure was 100 kPa. The mass spectrometer was used with an electron energy of 70 eV, the source at 240 °C, and an electron multiplier voltage of ~2000 eV. The injection port was kept at 300 °C. The sample was injected in splitless mode and the injection port was purged 2 min after the injection. The oven temperature program started at 100 °C, which was held for 1 min, increased at 30 °C/min to 160 °C, and then increased at 10 °C/min to 310 °C, which was held for 10 min. The solvent delay was 4 min, then selected ion monitoring mode was used. The m/z ratios 342, 432, and 213 were monitored to allow UV-327, UV-234, and BP-12, respectively, to be determined. The m/z ratios for the surrogate standards and internal injection standard were m/z 359 (UV-327-d20), m/z 343 (BP-12-d17), and m/z 240 (chrysene-d12). Quantification of each UVS was achieved by comparing the integrated peak area of the relevant quantification ion with the integrated peak area of the internal injection standard and using the relevant calibration line.

The BFRs were determined using a 7890 gas chromatograph with a micro electron capture detector (Agilent Technologies). Separation was achieved using a DB-5 15 m fused silica capillary column (0.25 mm i.d., 0.25 mm film thickness; Agilent Technologies). The carrier gas was helium. BDE-208 and f-BDE-208 were clearly separated and the BDE-209 and DBDPE peak heights maximized using ramped pressure mode. The column pressure was kept at 20.19 psi for 31 min, increased at 15 psi/min to 35 psi, and then stayed at 35 psi for 6 min. The injection port was kept at 250 °C. The sample was injected in splitless mode, then the injection port was purged 0.75 min after the injection. The oven temperature program started at 80 °C, which was held for 2 min, increased at 30 °C/min to 240 °C, and then increased at 3 °C/min to 300 °C, which was held for 15 min. The BFRs were identified and quantified against the relevant standards.

#### Analytical quality control and assurance

A procedural blank was analyzed with every set of samples. An analyte concentration less than three times the concentration found for the blank samples was classed as below the limit of quantification. The quality control and assurance procedures included analyzing four replicate mussel samples that had been spiked with the target compounds. The relative standard deviations of the BDE-209, DBDPE, UV-327, UV-234, and BP-12 concentrations were 2%, 2%, 2%, 10%, and 12%, respectively. The mean BDE-209, DBDPE, UV-327, UV-234, and BP-12 recoveries were 90%, 114%, 92%, 100%, and 83%, respectively.

### Statistical Analysis

Statistical analyses were performed using R [17]. Differences between the amounts of additives accumulated in the mysid, fish muscle, and fish liver samples from different treatments were assessed using generalized linear models with log link functions. A gamma distribution was assumed to account for the positive continuous values because Shapiro–Wilk normality tests indicated that the response variables had non-normal distributions. The log-transformed sample dry weight was used as an offset in each model. Effects of the treatments were assessed by performing likelihood ratio tests using the Anova function in the “car” package [18]. Pairwise comparisons were performed using Tukey’s honestly significant difference tests for post-hoc analysis.

## Results

### Leaching experiment

Overall, 2.33% of the additives were leached from the polyethylene microplastics in 24 h. Different leaching rates were found for the different additives (Table 2). The mean leaching rate was 3.16%. BP-12 had a higher leaching rate (8.79%) than the other additives (<3%).

**Table 2.** Results of the leaching experiment performed using 2 mg of microplastics in 1 L of seawater for 24 h.

| Additive | Blank | Initial amount<br>in plastics (ng) |  | Final amount<br>in seawater (ng) |  | Leaching rate<br>(%) |  |
| --- | --- | --- | --- | --- | --- | --- | --- |
|  |  | Mean | SE | Mean | SE | Mean | SE |
| BDE 209 | 0.127 | 713 | 60951 | 6.27 | 5.27 | 0.86% | 0.72% |
| DBDPE | 60.0 | 148 | 21930 | 4.13 | 3.53 | 2.61% | 2.20% |
| UV 327 | 0.208 | 722 | 66262 | 5.12 | 1.34 | 0.70% | 0.18% |
| UV 234 | 0.327 | 794 | 9662 | 23.2 | 5.94 | 2.83% | 0.71% |
| BP 12 | 4.83 | 246 | 12082 | 23.8 | 3.14 | 8.79% | 1.07% |

SE is the standard error. Gray cells indicate concentrations below the limits of quantification. All of the DBDPE concentrations were technically below the limit of quantification because the blank concentration was very high.

### Ingestion experiment using mysids

The concentrations of the five additives in the mysids from the different treatments were significantly different (Fig 1). The additive concentrations were 4–2450 times higher in the mysids that had ingested microplastics than in mysids analyzed straight after being collected from the ambient environment. The additive concentrations were slightly higher in the mysids treated with leachate seawater than in mysids analyzed straight after being collected from the ambient environment. The BP-12 concentrations in mysids treated with leachate seawater and mysids exposed to microplastics were not significantly different.

**Fig 1.**
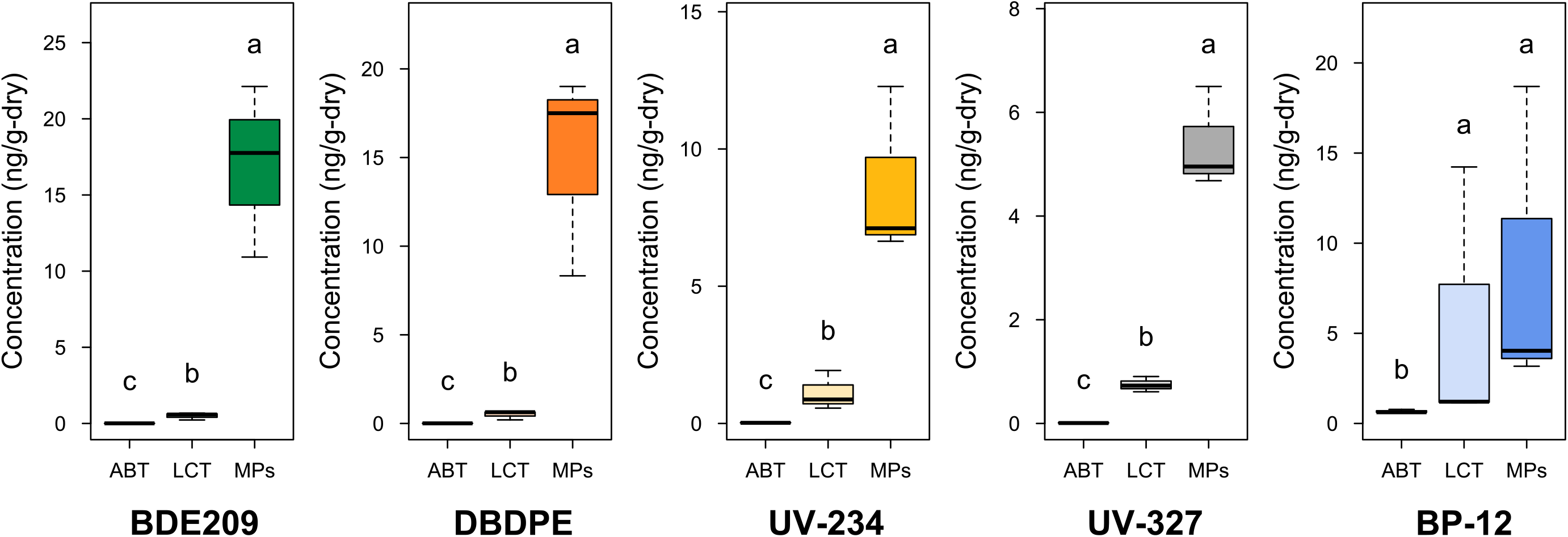
Concentrations of the five additives in individual mysids collected from the ambient environment (ABT), treated with leachate seawater (LCT), and exposed to microplastics (MPs). The median is shown as a solid horizontal line within a box, the 25th and 75th percentiles are the bottom and top of the box, respectively, and the 10th and 90th percentiles are the whiskers. Different letters indicate significant differences identified by performing post-hoc comparisons (p < 0.05 generalized linear model with a post-hoc Tukey’s honestly significant difference test).

### Trophic transfer experiment

The concentrations of the five additives were significantly different in the tissues from the fish directly and indirectly exposed to microplastics but the magnitudes of the differences in the concentrations were very different for the different additives and tissues, as shown in Fig 2. The BFR concentrations were higher in muscle from fish that were fed plastic-exposed mysids than in muscle from fish exposed to microplastics suspended in the water column. In contrast, the BDE-209 and DBDPE concentrations in liver from fish that were fed mysids and exposed to microplastics suspended in the water column were similar but more than three times higher than the concentrations in liver from the ambient control fish.

**Fig 2.**
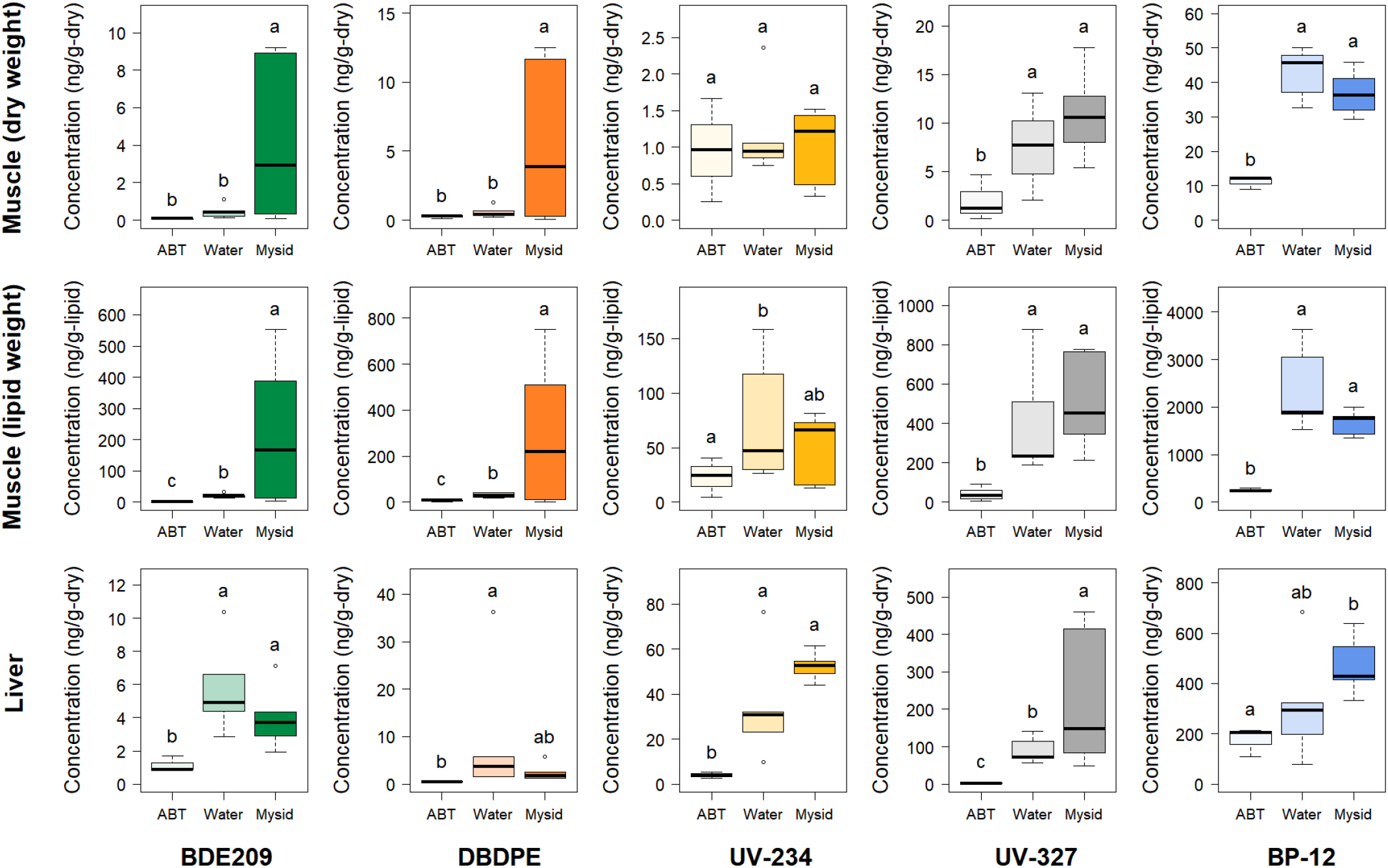
Concentrations of the five additives in muscle (dry-weight-basis and lipid-weight-basis) and liver (dry-weight-basis only) from fish immediately after collection from the ambient environment (ABT), fish exposed to microplastics suspended in the water column (Water), and fish fed mysids previously exposed to microplastics (Mysid). In each box and whisker plot, the median is shown as a solid horizontal line, the interquartile range (25th to 75th percentiles) is shown by the lower and upper ends of the box, and the 10th and 90th percentiles are shown as whiskers. Different letters indicate significant differences identified by performing post-hoc comparisons (p < 0.01 for a generalized linear model with post-hoc Tukey’s honestly significant difference tests).

The UV-327 and BP-12 concentrations in muscle were three–five times higher in the muscle samples from the treatment groups than in the muscle samples from the ambient controls. Unlike for BDE-209 and DBDPE, however, the UV-327 and BP-12 concentrations in muscle from fish fed mysids and exposed to microplastics in the water were not significantly different. The UV-234 and UV-327 concentrations were 9–13 and 35–89 times higher, respectively, in liver from fish in the treatment groups than the ambient controls, and the UV-327 concentration was significantly higher in liver from fish fed mysids than from fish exposed to microplastics in the water. The BP-12 concentration was three times higher in liver from fish fed mysids than the ambient control fish but the BP-12 concentration in liver from fish exposed to microplastics in the water was not significantly different from the concentration in liver from fish fed mysids and the ambient control fish.

The additive concentrations in the fish muscle samples were also calculated on a lipid weight basis. Similar accumulation patterns were found for the additives on a lipid weight basis as for the additives on a dry weight basis. The concentrations of both BFRs were significantly higher in the samples from the fish fed mysids that had been exposed to microplastics than in the samples from the fish exposed to plastic-contaminated water. However, the mean effects were markedly stronger for the lipid-weight-basis results (factors of 161 and 52 for BDE-209 and DBDPE, respectively) than for the dry-weight-basis results (factors of 71 and 22 for BDE-209 and DBDPE, respectively). Likewise, UVSs accumulation in the different treatments followed similar patterns using the lipid-weight-basis and dry-weight-basis results. The mean UVSs concentrations were higher in the samples from the fish fed mysids that had been exposed to microplastics than in the samples from the fish exposed to plastic-contaminated water by factors of 7–12 on a lipid weight basis and by factors of 5–7 on a dry weight basis.

## Discussion

It has been found that microplastics can act as vectors of plastic-associated chemicals to marine organisms, but most previous studies have focused on environmental contaminants adsorbed to microplastics rather than contaminants added to the microplastics during manufacture [19,20]. Additives to plastics are different from pollutants that adsorb to plastics in that additives are not attached to the polymer surfaces but are incorporated at high concentrations into the plastics [11]. Positive relationships between plastic ingestion and chemical additive concentrations in marine fish and seabird tissues have been found in field studies [21,22], suggesting that plastic-derived additives can be transferred to the tissues of organisms in natural environments. Our results support this, as do the results of a previous laboratory study in which plastic ingestion led to additives accumulating in the tissues of seabirds [8]. Fluids in the guts of seabirds and fish have been found to leach chemicals from plastics [9,23], which would explain chemical additives accumulating in the tissues of organisms after plastic has been ingested. Plastic ingestion has been reported in > 363 of fish species [1], so our results indicate that there should be concern about the exposure of fish to chemicals that are added to plastics.

We found that BDE-209 and DBDPE accumulation occurred in fish more strongly through prey-mediated ingestion of microplastics than through ingestion of waterborne microplastics. There are two possible explanations for this. First, a fish could be exposed to larger amounts of chemical additives through ingesting mysids exposed to microplastics than through directly ingesting microplastics. We previously found that fish ingested 3–11 times more microplastics in mysids than from the water column [4]. A higher microplastic ingestion rate through indirect ingestion than direct ingestion would result in more exposure to chemical additives in the fish stomach through indirect ingestion than direct ingestion. Second, fragmentation of microplastics by mysids could facilitate chemicals being released from the microplastics. Some crustacean species, including mysids [4,24], can fragment plastics into smaller particles. Chemicals will generally be released more readily by smaller than larger plastic particles because the surface-to-volume ratio will be higher for smaller than larger particles [25]. Ingesting microplastics in mysids, therefore, have affected the sizes of the microplastic particles in the fish stomach and may have enhanced leaching of the chemicals in the microplastic particles.

Although we found larger amounts of BDE-209 and DBDPE accumulated in fish tissues through prey-mediated exposure to microplastics than through direct exposure to microplastics, direct exposure and prey-mediated exposure contributed equally to accumulation of UV-327, UV-234, and BP-12 in the fish tissues. UV-327, UV-234, and BP-12 are likely to leach from plastics more easily than BDE-209 and DBDPE because UV-327, UV-234, and BP-12 are less hydrophobic (log Kow 6.91–7.67) than BDE-209 and DBDPE (log Kow 12.11–3.64). In particular, the leaching rate for BP-12 was 8.79%, which was higher than the leaching rates of the other additives (Table 2). Surfactants added to disperse the polyethylene particles in the water could also have facilitated leaching. UV-327, UV-234, and BP-12 may be taken up more readily than BFRs directly from the water through fish gills. Although our results indicate that accumulation patterns of plastic additives vary depending on the properties of the chemicals, both direct and indirect pathways for microplastic ingestion are important for plastic-derived chemical accumulation in fish.

Microplastics have been found to be ingested by >701 species of marine organisms [1]. Various chemicals are added to almost all plastics, so most species of marine organisms are likely to be affected by additives to plastics. Organisms may be more seriously affected by organic compounds such as plastic additives than by plastics themselves because the additives may be persistent in the tissues and have physiological effects. Our results indicated that additives to plastics can be transferred through the food chain along with microplastics and raise concerns about the risks posed by additives to plastics to organisms at high trophic levels, including humans. Sustainable marine resource exploitation and food security will require more rigorous regulations and management of plastics and the chemicals added to plastics.

## Conclusion

This study is the first to demonstrate that trophic transfer of microplastics has a higher contribution to the accumulations of plastic additives (BDE209 and DBDPE) in fish tissues than direct ingestion from the water column. However, as some chemicals (UV-234, UV-327, and BP-12) showed similar accumulation patterns between the two exposure pathways (direct ingestion from the water column vs. indirect ingestion via trophic transfer), their relative contributions vary among types of additives. Therefore, both direct and indirect exposure pathways are important in accumulations of plastic additives in marine fish. As many additives are potentially toxic, this study raises the concern about their negative impacts on organisms at high trophic levels. To better understand the fates and the effects of plastic additives accumulated in organisms, further research should focus on their in vivo kinetics and toxicity.

## Acknowledgments

We gratefully thank Captain S. Hamano, H. Katsuragawa, and other staff of the Akkeshi Marine Station for assistance with sample collection. We thank Gareth Thomas, PhD, from Edanz (https://jp.edanz.com/ac) for editing a draft of this manuscript.

## Supporting Information

**Fig S1. Size distribution of polyethylene microplastics used in the exposure experiment.**

The red line is the mean particle size.

**Fig S2. Contributions of the additives to the total additive concentration in the polyethylene microplastics used in the experiment.**

**Table S1. Results of likelihood ratio tests performed on the generalized linear models of the concentrations of the additives in mysids as a function of the treatment (ambient control, leachate treatment, and microplastic treatment).**

**Table S2. Results of likelihood ratio tests performed on the generalized linear models of the concentrations of the additives in fish muscle and liver as a function of the treatment (ambient control, water treatment, and mysid treatment).**

**Dataset S1. Concentrations of the target additives in the mysid and fish samples collected after the exposure experiments.**

## Notes

### Competing Interest Statement

The authors have declared no competing interest.

### Summary of Updates

Introduction and Discussion were revised to be more concise. Information on microplastics and plastic additives, together with results of likelihood ratio tests, were included in the supplementary information.

