## Supplementary Information for "Trophic transfer of microplastics enhances plastic additive accumulation in fish"

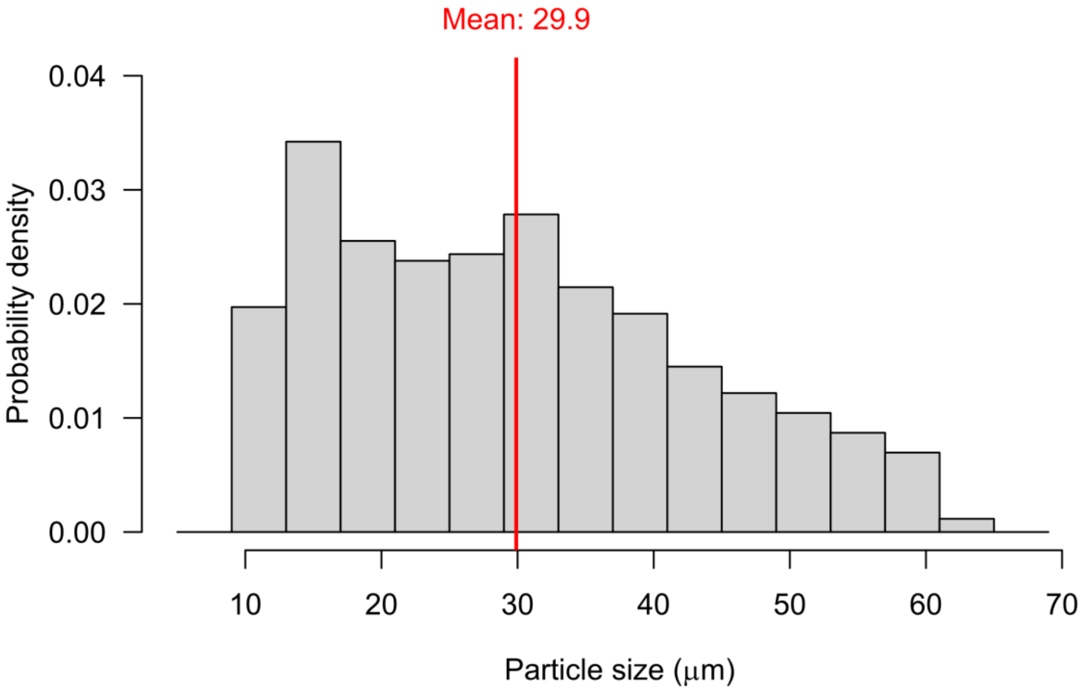


**Fig S1. Size distribution of polyethylene microplastics used in the exposure experiment.** The red line is the mean particle size.

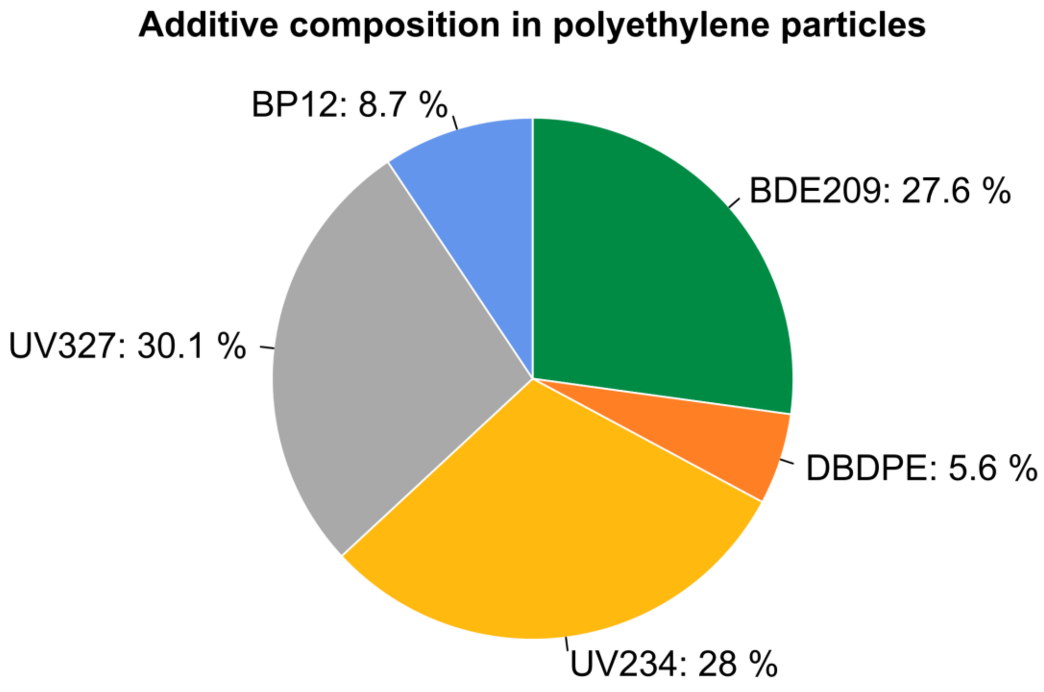


**Fig S2. Contributions of the additives to the total additive concentration in the polyethylene microplastics used in the experiment.**

**Table S1. Results of likelihood ratio tests performed on the generalized linear models of the concentrations of the additives in mysids as a function of the treatment (ambient control, leachate treatment, and microplastic treatment).**

| Additive | Fixed factor | LR Chisq | df | *p*-value |
|---|---|---|---|---|
| BDE209 | Treatment | 253 | 2 | < 0.001 |
| DBDPE | Treatment | 186 | 2 | < 0.001 |
| UV234 | Treatment | 134 | 2 | < 0.001 |
| UV327 | Treatment | 280 | 2 | < 0.001 |
| BP12 | Treatment | 10.8 | 2 | 0.004 |

**Table S2.**

Results of likelihood ratio tests performed on the generalized linear models of the concentrations of the additives in fish muscle and liver as a function of the treatment (ambient control, water treatment, and mysid treatment).

| **Body tissue** | **Additive** | **Fixed factor** | **LR Chisq** | **df** | ***p*-value** |
|---|---|---|---|---|---|
| Muscle (Dry weight) | BDE209 | Treatment | 27.9 | 2 | < 0.001 |
| | DBDPE | Treatment | 27.2 | 2 | < 0.001 |
| | UV234 | Treatment | 0.441 | 2 | 0.802 |
| | UV327 | Treatment | 9.72 | 2 | 0.008 |
| | BP12 | Treatment | 99.5 | 2 | < 0.001 |
| Muscle (Lipid weight) | BDE209 | Treatment | 38.2 | 2 | < 0.001 |
| | DBDPE | Treatment | 36.4 | 2 | < 0.001 |
| | UV234 | Treatment | 5.15 | 2 | 0.076 |
| | UV327 | Treatment | 17.6 | 2 | < 0.001 |
| | BP12 | Treatment | 91.0 | 2 | < 0.001 |
| Liver (Dry weight) | BDE209 | Treatment | 18.5 | 2 | < 0.001 |
| | DBDPE | Treatment | 11.9 | 2 | 0.003 |
| | UV234 | Treatment | 33.4 | 2 | < 0.001 |
| | UV327 | Treatment | 40.4 | 2 | < 0.001 |
| | BP12 | Treatment | 7.01 | 2 | 0.030 |
